# Physiological constraints on the rapid dopaminergic modulation of striatal reward activity

**DOI:** 10.1101/2022.09.16.508310

**Authors:** Charltien Long, Kwang Lee, Long Yang, Theresia Dafalias, Alexander K. Wu, Sotiris C. Masmanidis

## Abstract

While the contribution of dopaminergic (DA) neurons to associative learning is firmly established, their importance for influencing imminent behavior on short (subsecond) timescales is less clear. Mechanistically, it is thought that DA neurons drive these behavioral changes because of their ability to rapidly alter striatal spiking activity. However, due to limitations of previous approaches, the straightforward prediction that striatal spiking is rapidly influenced by physiologically relevant DA signals has not been rigorously tested. Here, we monitored changes in spiking responses in the ventral striatum while transiently reducing or increasing DA levels. Contrary to the predicted effect, neither spontaneous nor reward-evoked striatal spiking activity was strongly influenced by optogenetic manipulations, except when DA exceeded reward-matched levels. These findings challenge the view that DA plays a major role in rapidly influencing striatal activity. Finally, they suggest a need to distinguish between the modulatory functions of DA under physiological and supra-physiological conditions.

## Introduction

The striatal regulation of learning, movement, motivation, and decision-making is thought to critically depend on DA signaling ^1^. Abnormal levels of striatal DA are implicated in several disorders including Parkinson’s disease, addiction, and depression ^2, 3^. Yet despite significant progress, our understanding of DA’s complex modulatory functions in the striatum is incomplete, and remains the subject of intense study and controversy. Nevertheless, it is commonly accepted as dogma that striatal DA serves two major functions ^4^. First, it regulates synaptic plasticity, a protracted process that is necessary for associative learning ^5, 6^. Second, under certain conditions DA neurons have been found capable of strongly influencing striatal neural spiking activity on short, subsecond timescales ^7, 8^. Rather than contributing to learning, this second, rapid modulatory process appears better suited for influencing imminent or ongoing behaviors. This view has earned wide support from behavioral studies suggesting that DA plays a major role in online movement initiation, reward-motivated approach, and action selection ^9–14^. This appears further buttressed by data that, in addition to canonical reward prediction error (RPE) signals ^15^, DA neurons and their striatal projections encode information about rapid behavioral processes such as movement initiation ^16–18^.

Despite this evidence, an opposing view has emerged that DA only plays a minor role in subsecond behavioral control, and is primarily involved in associative learning ^19–21^. But crucially, this role can be restored when DA levels are artificially raised above physiological levels ^12, 19^. Since this supra-physiological regime is thought to reflect an atypical operating mode of DA circuits, if not properly identified as such, it may lead to a different interpretation of DA’s functional role.

The emergence of these two countervailing views suggests a pressing need to reconsider the direct electrophysiological evidence on which this argument hinges - that striatal spiking activity is strongly and rapidly influenced by physiological DA signals. On one hand, there is a large body of literature using on in vivo and ex vivo measurements to show that DA indeed influences striatal activity. Early efforts to understand DA’s modulatory effects on striatal activity used iontophoretically applied DA ^22–24^, which was not calibrated to physiological DA levels, or electrical stimulation of the midbrain or medial forebrain bundle, which non-specifically activated DA circuits ^25–27^. This was followed by pharmacological manipulation of DA neurons or receptors ^28, 29^, as well as chronic DA lesions ^30, 31^. These approaches are primarily intended to examine the protracted modulatory contributions of DA (e.g., synaptic plasticity) that occur over multi-second and longer timescales. In contrast, they are less suited for identifying rapid electrophysiological effects. In contrast, optogenetic manipulations address both the need for cellular and temporal specificity ^3, 7, 8^. However, previous studies based on this technique often did not take into account the potential distinction between physiological and supra-physiological DA levels. Consequently, while the presentation of food rewards raises striatal DA levels by 10-100 nM ^10^, numerous studies employing optogenetically evoked striatal DA release exceeded those levels. Thus, owing to a range of factors, there is a lack of conclusive electrophysiological evidence that reward-matched or other physiologically calibrated DA signals play a major role in controlling striatal spiking patterns on subsecond timescales.

Here we examined the ability of DA neurons to elicit rapid changes in striatal spiking activity. We focused on the role of DA in the ventral striatum during the delivery of unexpected rewarding stimuli, because these stimuli elicit large and rapid DA responses in the ventral striatum ^32^. Since unexpected rewards induce concurrent changes in striatal firing patterns ^33^, we tested whether DA signals are necessary or sufficient for driving reward-evoked neural activity in the ventral striatum. Experiments were carried out by optogenetically subtracting or adding DA activity while simultaneously monitoring ventral striatal DA and electrophysiological activity. This combined approach enabled a systematic study of DA’s rapid modulatory effects under reward-matched and supra-reward signaling conditions.

## Results

### Simultaneous monitoring of DA and electrophysiological activity in the ventral striatum

In order to measure neural population activity in the ventral striatum under calibrated levels of change in DA signaling, we constructed an opto-probe device consisting of a multielectrode recording array on a silicon microprobe, coupled to an optical fiber for photometry (**Fig. 1a**). The optical fiber was used for fluorescence-based DA monitoring in the vicinity of the electrodes with the genetically encoded sensor dLight1.2 ^34^. The opto-probe was implanted in the nucleus accumbens area of the ventral striatum in head-restrained food-restricted mice. In each animal, the electrodes captured the spiking response of tens of single-units (mean ± SD: 113 ± 30 neurons) alongside photometric data on local DA signaling changes. Another optical fiber was implanted in the ventral tegmental area (VTA) to allow for optogenetic manipulation of DA neurons using virally mediated Cre-dependent opsin expression in DAT-Cre mice (**Fig. 1b**). Unconditional rewarding stimuli (sweetened milk) were delivered at random intervals to elicit robust increases in licking and DA release (**Fig. 1c**). Concurrently, the majority (65 %) of recorded neurons in the ventral striatum were modulated by unexpected rewards (**Fig. 1d,e**) ^33^. Neurons showed a predominantly excitatory subsecond-scale initial reward response, which was sometimes followed by a more prolonged period of inhibition. A strong reward response was observed in all three major putative classes of electrophysiologically identified cell types, corresponding to medium spiny projection neurons (MSNs), fast spiking interneurons (FSIs), and tonically active projection neurons (TANs) (**Supplemental Fig. 1**). However, to avoid possible bias arising from cell-type specific modulatory effects or classification errors, all subsequent analysis used spiking data from every recorded neuron.

**Fig. 1:**
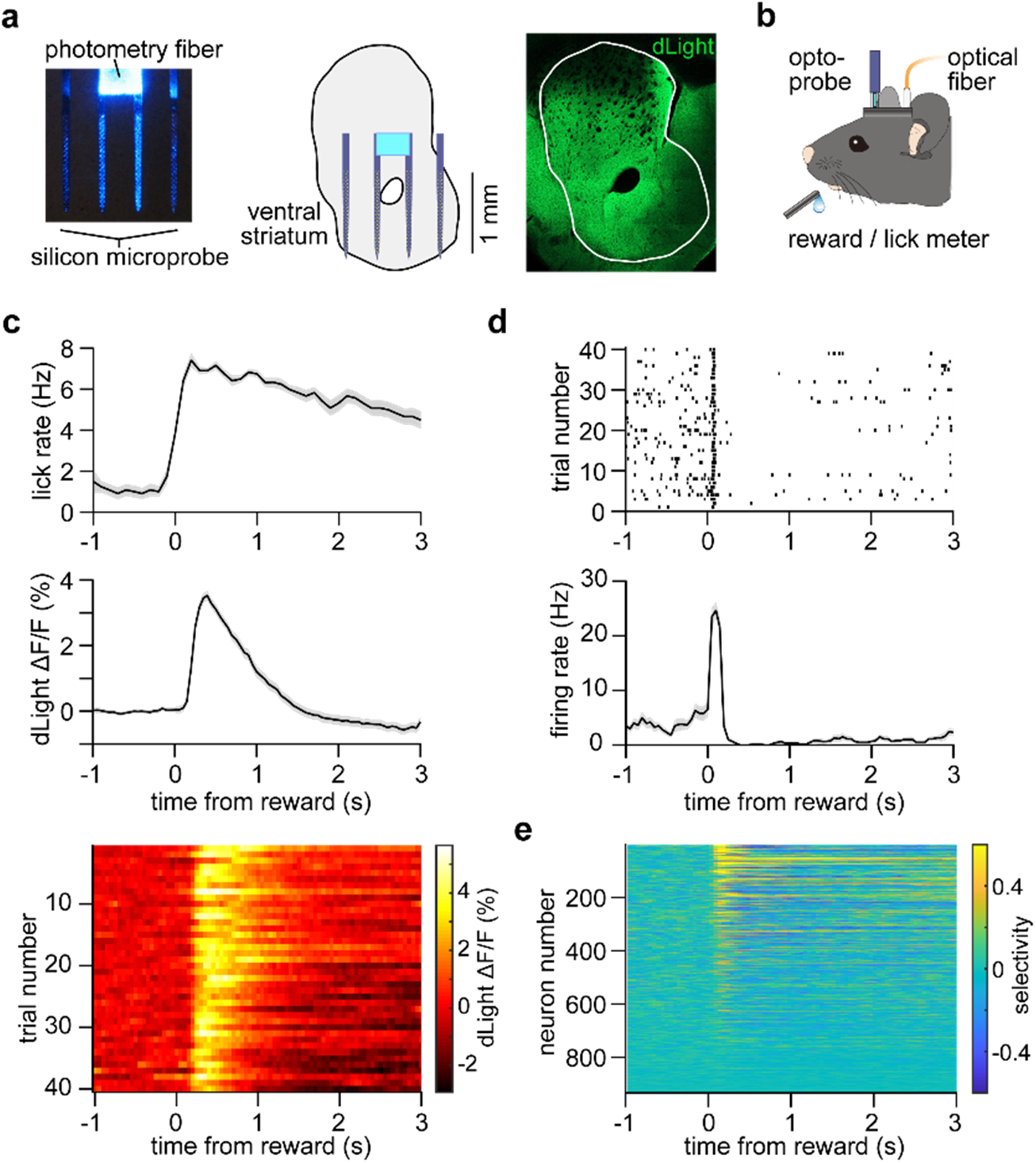
Simultaneous monitoring of DA and electrophysiological activity in the ventral striatum. a. Left: Opto-probe device comprised of a 256 electrode array distributed on four shanks, together with an optical fiber for photometry. Middle: Illustration of the opto-microprobe in the targeted area of the ventral striatum. Right: Confocal image of dLight1.2 expression. b. Mouse recording apparatus during reward delivery and optogenetic stimulation. c. Reward-evoked licking and dLight fractional fluorescence change data from one animal. Spike raster and mean firing rate of one reward-responsive MSN. d. Selectivity index of 933 ventral striatal neurons during reward delivery. Selectivity of neural activity is calculated relative to baseline. All data in this figure represent mean ± SEM.

These combined measurements revealed temporally correlated changes in ventral striatal DA and spiking activity during reward delivery. Based on the view that DA neurons drive changes in spiking on subsecond timescales, it has been posited that these two forms of reward signals are causally related ^27, 35^. We therefore manipulated DA activity in order to directly test this prediction.

### Small effect of inhibiting DA reward transients on striatal spiking activity

First, we used opto-inhibition to examine whether DA neuron signaling is necessary for reward-evoked striatal spiking activity (**Fig. 2a**). We expressed eNpHR3.0 in VTA DA neurons, and timed laser stimuli to occur together on 50 % of reward trials (5 mW, 0.5 s continuous laser). Trials consisting of only reward (R) and laser-paired reward (R+L) were randomly interleaved. As expected, reward-evoked DA release in eNpHR3.0 expressing animals, but not opsin-free controls, was significantly inhibited and even transiently fell below baseline levels (**Fig. 2b,c**). The duration of opto-stimulation was intentionally matched to previous studies supporting the involvement of DA neurons in rapid behavioral control ^10, 11^. Consummatory licking was not significantly altered by DA neuron inhibition, which was applied unilaterally (**Fig. 2d**). This allowed us to examine changes in neural dynamics in the absence of potentially obfuscating behavioral changes ^29^. According to a prominent view of DA function, transiently reducing DA levels should cause strong and rapid changes in reward-evoked spiking responses in the ipsilateral ventral striatum. To test this we directly compared differences in individual neuron firing profiles between reward and laser-paired reward trials (**Fig. 2e,f**). These differences were quantified with a selectivity index based on the area under the receiver operating characteristic (ROC) curve (range of index: ±1). Positive/negative index values denote higher/lower spike rates during DA inhibition, respectively. We also applied a statistical test to identify bins with significant differences in firing (see Methods). In contrast to the pronounced change in DA signals, concurrent changes in spiking activity were considerably more subtle (**Fig. 2g**). DA neuron inhibition altered the reward response of 11 % of neurons, compared to 10 % in control animals – a statistically insignificant difference (**Fig. 2h**). On average the magnitude of the selectivity index also differed from controls by a small though statistically significant amount (**Fig. 2i**).

**Fig. 2:**
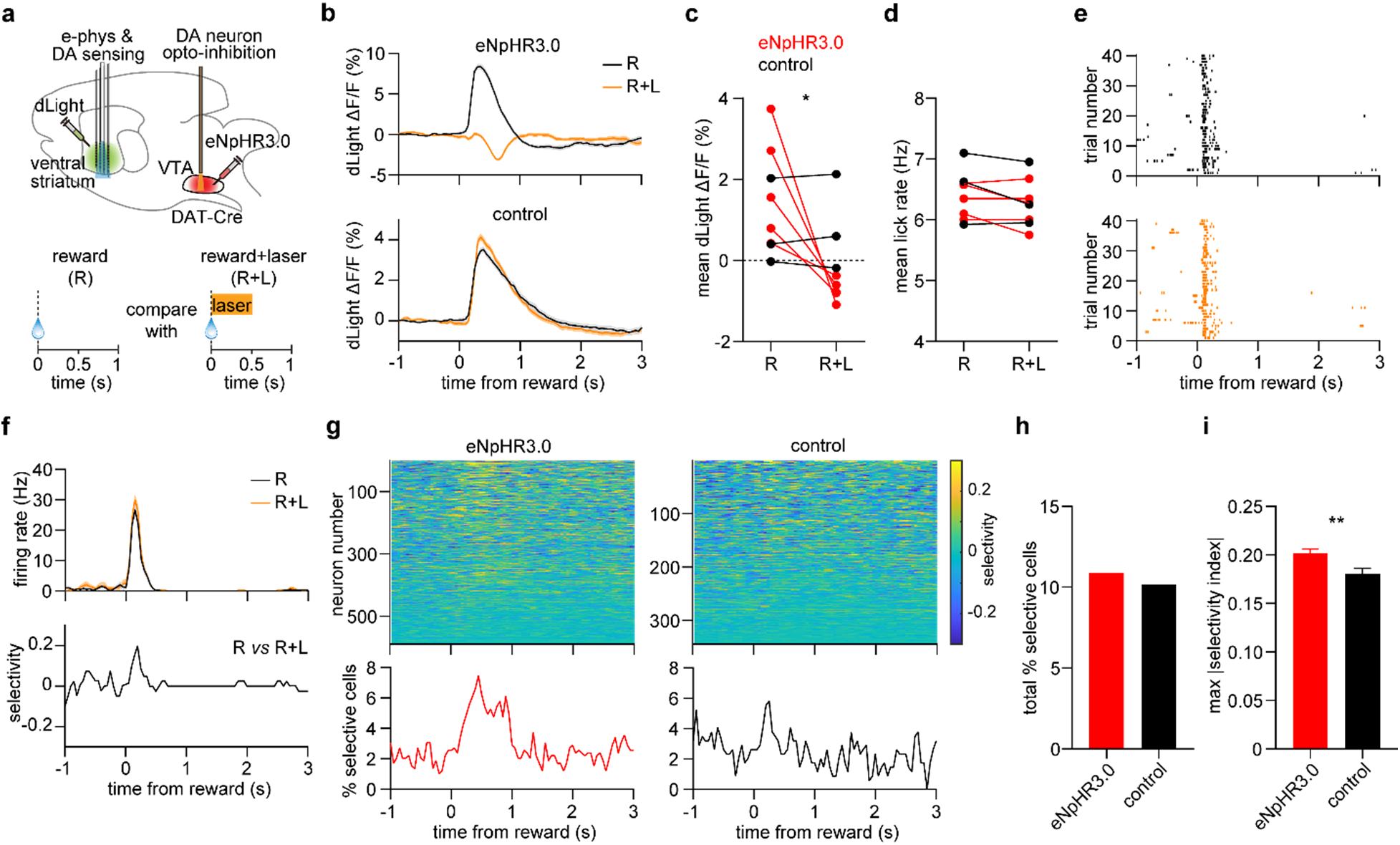
Small effect of inhibiting VTA DA neurons on reward-evoked striatal spiking activity. a. Top: Experimental approach. Bottom: task schematic in which reward trials (R) are compared to laser-paired reward trials (R+L). b. dLight fractional fluorescence change signal from one eNpHR3.0-expressing mouse (top) and one opsin-free control mouse (bottom). c. Mean reward-evoked dLight fluorescence on R and R+L trials (n = 5 eNpHR3.0 and 3 control mice, two-way RM ANOVA, group effect: *F_1,6_* = 0.2, *P* = 0.7, trial effect: *F_1,6_* = 7, *P* = 0.04). d. Mean reward-evoked lick rate on R and R+L trials (n = 5 eNpHR3.0 and 3 control mice, twoway RM ANOVA, group effect: *F_1,6_* = 0.5, *P* = 0.5, trial effect: *F_1,6_* = 3.8, *P* = 0.1). e. Spike raster of one reward-responsive neuron on R (top) and R+L (bottom) trials. Data are from an eNpHR3.0-expressing animal. f. Mean firing rate (top) and selectivity index (bottom) of the same neuron as panel e. Selectivity is calculated between R and R+L trials. g. Top: Selectivity index of 589 neurons pooled across 5 eNpHR3.0-expressing mice, and 344 neurons pooled across 3 control mice. Bottom: Percentage of neurons that were significantly selective for R versus R+L trials, as a function of time. h. Total percentage of neurons that were selective for R versus R+L trials (n = 64 out of 589 cells in the eNpHR3.0 group, n = 35 out of 344 cells in the control group, chi square test for proportions, *χ^2^* = 0.2, *P* = 0.8). i. Maximum absolute value of the selectivity index per neuron (n = 589 cells in the eNpHR3.0 group, n = 344 cells in the control group, unpaired t-test, *t* = 3.1, \*\**P* = 0.002). All data in this figure represent mean ± SEM.

The above results reveal a small but statistically significant effect on individual neuron firing properties. It is still conceivable that these subtle electrophysiological changes may be important for regulating striatal dynamics and, in turn, behavior, on subsecond timescales. Indeed, if DA inhibition weakly but consistently alters the activity pattern of even a small group of striatal neurons, in principle, downstream brain areas could reliably read out this information to guide behavior. To test this possibility, we attempted to distinguish population-level dynamics observed between reward and laser-paired reward trials (i.e., R versus R+L), using a machine learningbased decoder. Decoding accuracy, reflecting the percentage of correctly classified trials, was briefly elevated for following reward delivery, but similar size effects were found with data from opsin-free control mice (**Fig. 3a**). It is also notable that the decoder never performed significantly above chance levels, as defined by the 95^th^ percentile of decoders trained on trial-shuffled data. Decoder performance using data from eNpHR3.0-expressing mice improved by training the classifier with higher numbers of neurons, and eventually surpassed control data (**Fig. 3b**). However, even under the most optimal decoder training conditions using spiking data from 300 neurons, only an 8 % improvement in performance was observed (74 % accuracy from eNpHR3.0 group data, compared to 66 % from control group data). Thus, even with the aid of population-level recordings and machine learning tools, we could not consistently distinguish reward from laser-paired reward trials.

**Fig. 3:**
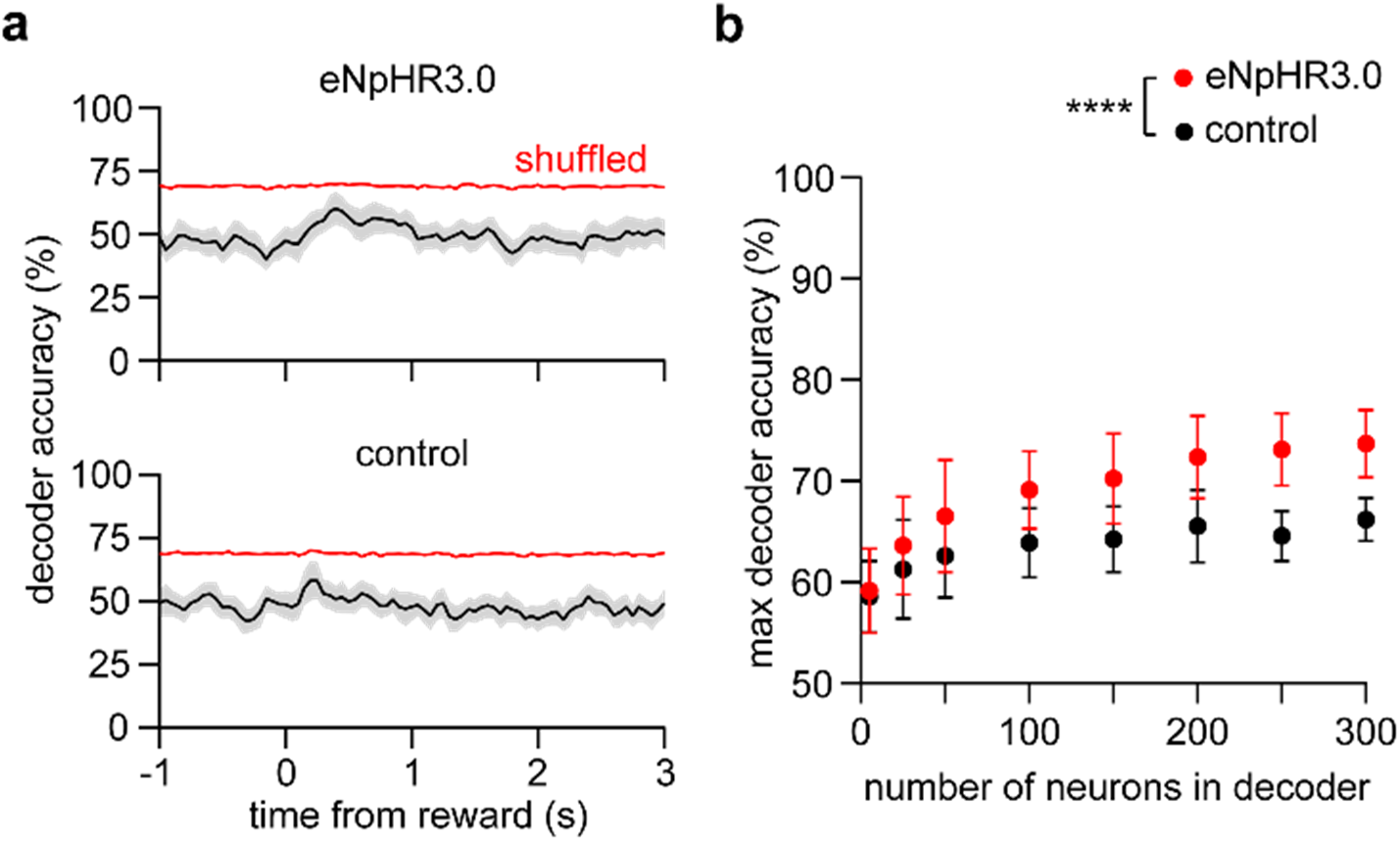
Population-level decoding discriminates striatal neuron reward responses with and without DA. a. Mean accuracy of an SVM decoder trained using 50 neurons to discriminate R from R+L trials. Red line indicates the 95 % confidence interval of decoder performance trained on trial-shuffled data. Top: neurons selected from the eNpHR3.0 group. Bottom: neurons selected from the control group. Shaded area represents represent the SD across 50 random drawings of neurons. b. Maximum decoder accuracy as a function of neuron number (two-way ANOVA, group effect: *F_1,784_* = 340, *P* < 0.0001, neuron number effect: *F_7,784_* = 92, *P* < 0.0001). Post-hoc Sidak’s test: \*\*\*\**P* < 0.0001 for all data points with n ≥ 50 neurons. Data represent the mean and SD across 50 random drawings of neurons.

On a subset of trials in the same experiment, we also examined DA contributions to spontaneous striatal spiking activity, by applying identical laser stimulation but without accompanying rewards (**Supplemental Fig. 2a**). This caused DA levels to transiently fall below baseline levels (**Supplemental Fig. 2b,c**). We looked for changes in neural activity relative to a baseline period preceding laser stimulation. Neither the fraction of selective cells nor the mean selectivity index were significantly different from controls (**Supplemental Fig. 2d-f**). A statistically significant difference arose when testing a decoder trained to distinguish laser-evoked population activity from baseline activity, but again the size of this effect was small (**Supplemental Fig. 2g,h**). Taken together, the results show that DA neuron inhibition has a weak effect on both reward-evoked and spontaneous spiking activity in the ventral striatum.

To ensure that the lack of strong electrophysiological effects cannot be attributed to problems with our recording or analysis approach, we carried out a control experiment involving activation of VTA GABA neurons, which locally regulate DA neuron firing and also project to areas including the ventral striatum (**Fig. 4a**). Since these neurons release an inhibitory neurotransmitter with rapid postsynaptic effects, we expected their contribution to striatal spiking activity would substantially exceed those of DA neurons. We expressed the excitatory opsin Chrimson in VTA GABA neurons, and delivered pulsed laser stimulation on a subset of reward trials. Activating these neurons produced a marked reduction in both reward-evoked striatal DA release and spiking activity, consistent with their inhibitory action (**Fig. 4b,c**). Similarly, at the population level, neural decoding performance reached 95 % accuracy with as few as 25 neurons used for training the classifier (**Fig. 4d**). VTA GABA neuron activation without reward delivery also had a large effect on spontaneous striatal firing rates (**Supplemental Fig. 3a-d**). These data indicate a stark contrast between the effect of VTA GABA neuron activation and DA neuron inhibition on striatal spiking activity.

**Fig. 4:**
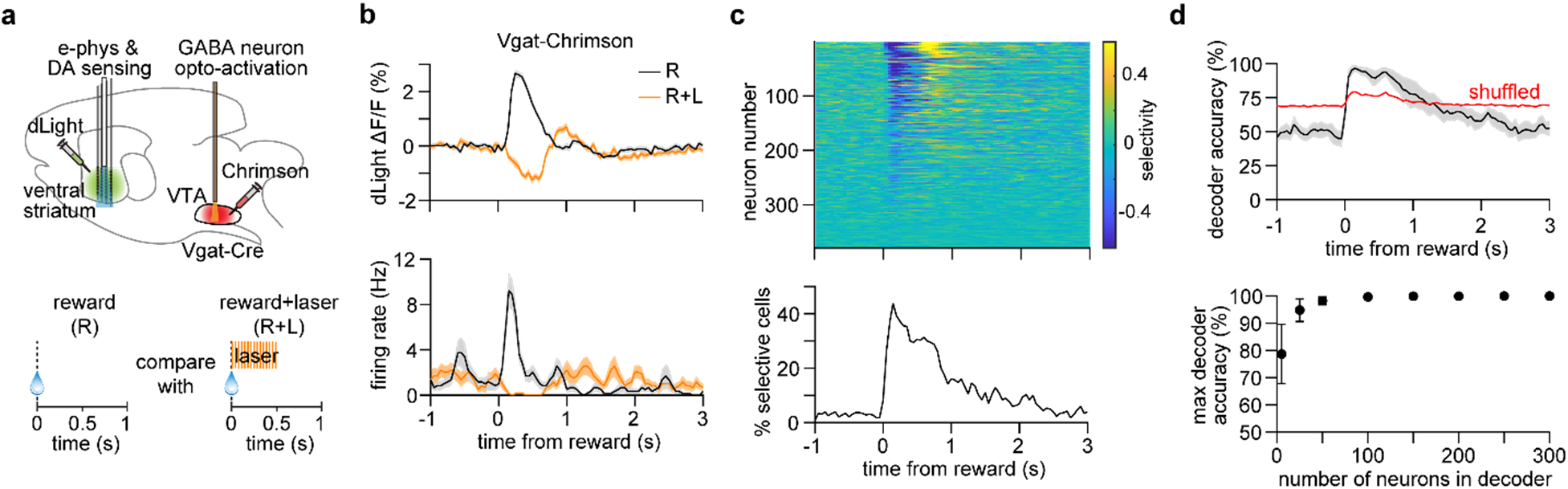
Strong effect of activating VTA GABAergic neurons on reward-evoked striatal spiking activity. a. Top: Experimental approach. Bottom: task schematic in which reward trials (R) are compared to laser-paired reward trials (R+L). Laser is pulsed at 40 Hz for 0.5 s total duration. b. Top: dLight fractional fluorescence change signal on R and R+L trials from one Vgat-Chrimson mouse. Bottom: Mean firing rate of one neuron on R and R+L trials. Data represent mean ± SEM. c. Top: Selectivity index of 379 neurons pooled across 3 Vgat-Chrimson mice. Bottom: Percentage of neurons that were significantly selective for R versus R+L trials, as a function of time. d. Top: Mean accuracy of an SVM decoder trained using 50 neurons to discriminate R from R+L trials. Shaded area represents the SD across 50 random drawings of neurons. Red line indicates the 95 % confidence interval of decoder performance trained on trial-shuffled data. Bottom: Maximum decoder accuracy as a function of neuron number (one-way ANOVA, neuron number effect: *F_7,392_* = 158, *P* < 0.0001). Post-hoc Sidak’s test: *P* < 0.0001. Data represent the mean and SD across 50 random drawings of neurons.

### Small effect of artificial reward-matched DA transients on spontaneous striatal activity

While our results suggest that DA signals during natural reward delivery only weakly contribute to striatal spiking activity, we cannot rule out the presence of enhanced effects under artificially elevated DA signaling regimes. To address this, we next used opto-activation to examine whether certain levels of DA are sufficient to produce strong and rapid changes in striatal spiking. The excitatory opsin Chrimson was expressed in VTA DA neurons, and pulsed laser stimulation was applied as before for a total duration of 0.5 s, both in isolation and paired with a subset of reward trials (**Fig. 5a**). To systematically study the effect of different DA levels on spontaneous striatal dynamics, we presented isolated laser stimuli and varied the stimulation frequency from 4 to 40 Hz in order to evoke increasing magnitudes of DA release (**Fig. 5b**). We then compared the peak level of optogenetically evoked to behaviorally evoked DA during reward delivery. The ratio between these values indicated the factor increase in DA above reward-matched levels. Across all sessions tested, this factor varied between 0.2 and 23, representing two orders of magnitude change in DA levels relative to reward. We next looked for laser-evoked changes in the spontaneous activity of simultaneously recorded neurons, relative to a pre-laser baseline period. Under certain conditions we found alterations in spontaneous firing upon DA neuron activation (**Fig. 5c**), but it was evident that the size of this effect depended on the level of DA release. Specifically, spontaneous spiking activity was only weakly sensitive to reward-matched DA levels, but appeared more strongly inhibited with higher, supra-reward DA levels (**Fig. 5d**). A similar trend emerged when testing a decoder trained to distinguish laser stimulation-evoked population activity from pre-laser baseline activity. Across all the data collected, we found an approximately linear change in the percentage of laser-modulated cells, selectivity index, and decoder performance as a function of the factor increase in DA (**Fig. 5e-g**). Together, these results suggest that artificial DA signals comparable to the magnitude of rewards are not on their own sufficient to cause large, rapid changes in striatal spiking. But at the same time, supra-reward DA transients are clearly capable of driving strong changes in neural activity.

**Fig. 5:**
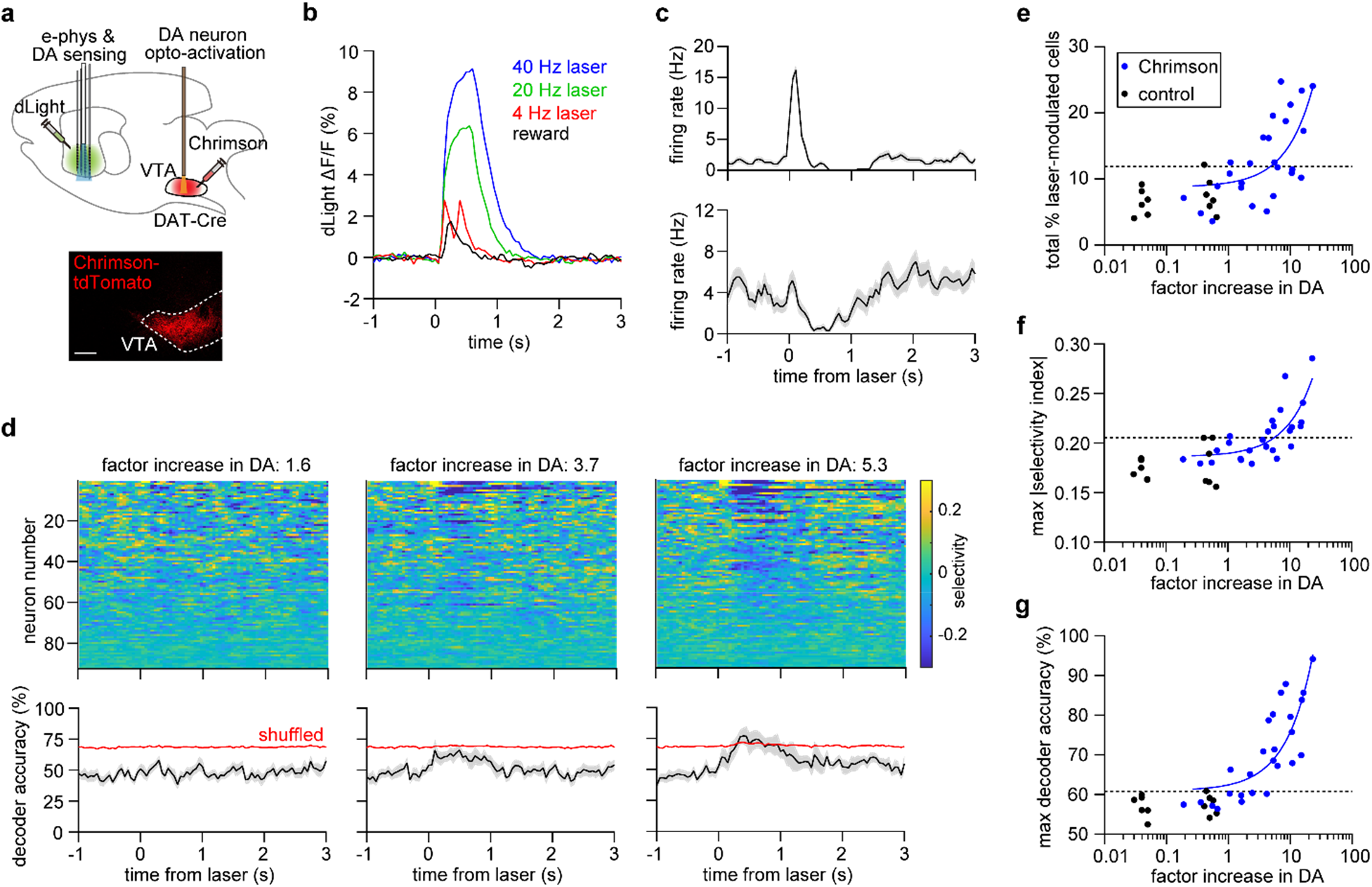
Small effect of artificial reward-matched DA transients on spontaneous striatal activity. a. Top: Experimental approach. Bottom: Confocal image of Chrimson-tdTomato expression in the VTA. b. dLight fractional fluorescence change signal from one Chrimson-expressing mouse. The data represent reward-evoked (black) and laser-evoked signals at 4 Hz (red), 20 Hz (green), and 40 Hz (blue) stimulation for 0.5 s. c. Mean firing rate of two laser-modulated MSNs. d. Top: Selectivity index of 92 simultaneously recorded neurons from one Chrimson-expressing mouse under 4 Hz (left), 20 Hz (middle), and 40 Hz (right) laser stimulation. Bottom: Mean accuracy of an SVM decoder trained using 50 neurons to discriminate laser-evoked from baseline activity. Shaded area represents the SD across 50 random drawings of neurons. Red line indicates the 95 % confidence interval of decoder performance trained on trial-shuffled data. e. Percentage of neurons per recording session that were significantly modulated by the laser, as a function of factor increase in DA. n = 26 sessions from 9 Chrimson-expressing mice, and n = 12 sessions from 4 control mice. Chrimson group data are positively correlated with the factor increase in DA (Pearson *r* = 0.6, *P* = 0.0006). Dashed line represents the 95 % confidence interval of the control data. Blue line represents the linear fit to the Chrimson data. f. Mean of the maximum absolute value of the selectivity index per recording session, as a function of factor increase in DA. Selectivity of neural activity is calculated relative to baseline. Chrimson group data are positively correlated with the factor increase in DA (Pearson *r* = 0.8, *P* < 0.0001 Dashed line represents the 95 % confidence interval of the control data. Blue line represents the linear fit to the Chrimson data. g. Maximum decoder accuracy per recording session, as a function of factor increase in DA. The decoder was trained using 50 neurons to discriminate laser-evoked from baseline activity. Chrimson group data are positively correlated with the factor increase in DA (Pearson *r* = 0.8, *P* < 0.0001). Dashed line represents the 95 % confidence interval of the control data. Blue line represents the linear fit to the Chrimson data.

Our estimate of the factor increase in DA may depend on several variables such as recording location. A large overestimate of this factor would be problematic, as it would weaken the conclusion that only supra-reward DA signals are sufficient to produce sizable electrophysiological effects. Spatial variations in dLight photometry signals are one potential source of uncertainty. Our opto-probe was configured such that the photometry fiber was placed above the most dorsal electrode recording sites, and thus was separated by over 1 mm from the most ventral recording sites (**Fig. 1a**). Since it is known that DA reward signals vary substantially across the striatum ^32^, the factor increase in DA cannot be assumed to be constant over the entire span of the electrode array along the dorsal to ventral axis. To quantify how this factor varies with depth, we performed a dLight photometry experiment with the optical fiber gradually moved more ventrally (6 different locations spanning 1 mm; **Supplemental Fig. 4a**). At each location we measured the magnitude of DA signals during reward delivery and optogenetic stimulation. While both of these signals varied considerably across different locations (**Supplemental Fig. 4b,c**), overall the factor increase in DA displayed a moderate increase with depth (**Supplemental Fig. 4d**). In experiments involving combined electrophysiology and photometry, access to DA signals from only near the most dorsal optical fiber position was available. Thus, these data suggest we may have underestimated the value of the factor increase in DA. If this is the case it would actually strengthen our conclusion. Another source of variability may relate to changes in DA reward signaling across successive trials. In particular, it is possible that the initial set of reward trials are more unexpected than latter trials, and therefore induce a stronger RPE response. To check if this impacts our estimate, we calculated a recalibrated factor increase in DA using the average of only the first 10 % of reward trials. On average the recalibrated factor was 17 % smaller than the original estimate, but this did not appear to fundamentally alter our conclusions (**Supplemental Fig. 4e,f**).

### Supra-reward DA signals modulate reward-evoked striatal spiking activity

Since the modulatory effects of DA are thought to be state-dependent ^1^, we also examined how amplifying DA reward signals alters ventral striatal spiking activity. Experiments involving DA neuron activation included a block of randomly interleaved reward and laser-paired reward trials. Pairing optogenetic stimulation with reward amplified the dLight fluorescence signal (**Fig. 6a,b**), but the consummatory licking response was not significantly altered (**Fig. 6c**). Consistent with our analysis of spontaneous activity, changes in reward-evoked neural activity were only clearly distinguished with high DA amplification (**Fig. 6d,e**). We next characterized how reward-evoked activity is altered under conditions when R and R+L trials could be reliably decoded (greater than 80 % decoder accuracy). In this supra-reward regime, neurons displayed a mixture of excitatory and inhibitory changes in firing (**Fig. 6f,g**). All three putative cell types exhibited changes in activity (**Supplemental Fig. 5**). MSNs showed mixed excitatory and inhibitory effects and the weakest overall changes, while the main influence on FSIs appeared to be inhibitory. A subset of TANs showed a broadened reward pause response, in qualitative agreement with previous work attributing the pause to DA modulation ^36^.

**Fig. 6:**
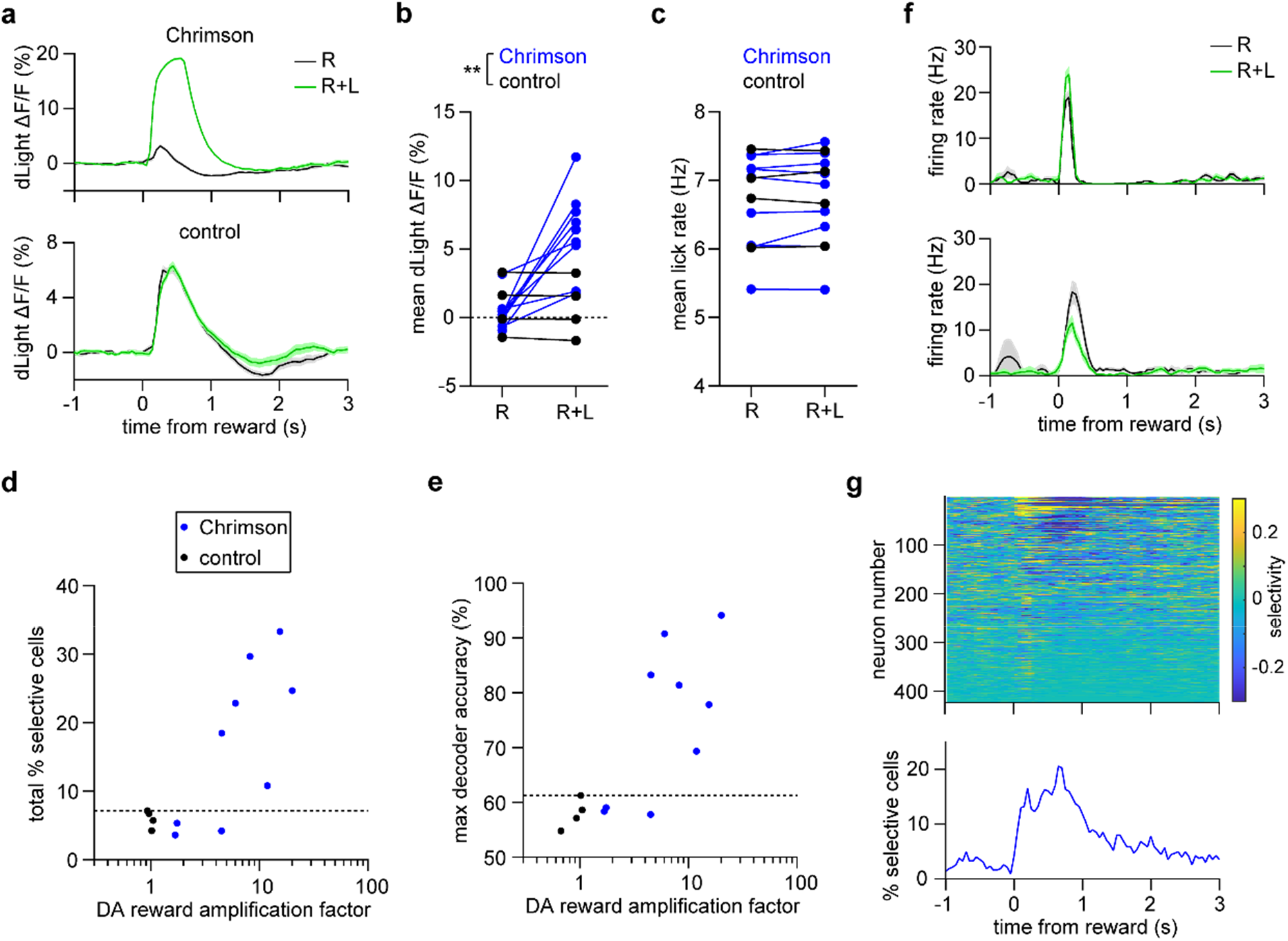
Supra-reward DA signals modulate reward-evoked striatal spiking activity. a. dLight fractional fluorescence change signal from one Chrimson-expressing mouse (top) and one opsin-free control mouse (bottom) on reward (R) and laser-paired reward (R+L) trials. Data represent mean ± SEM. b. Mean reward-evoked dLight fluorescence on R and R+L trials (n = 9 Chrimson and 4 control mice, two-way RM ANOVA, group effect: *F_1,11_* = 5.3, *P* = 0.04, trial effect: *F_1,11_* = 11, *P* = 0.007). Post-hoc Sidak’s test: \*\**P* = 0.001 for R+L trials. c. Mean reward-evoked lick rate on R and R+L trials (n = 9 Chrimson and 4 control mice, twoway RM ANOVA, group effect: *F_1,11_* = 0.1, *P* = 0.8, trial effect: *F_1,11_* = 0.6, *P* = 0.5). d. Percentage of neurons per animal that were selective for R versus R+L trials, as a function of DA reward amplification factor. n = 9 Chrimson-expressing and 4 control mice. Chrimson group data show a trend for being positively correlated with the DA amplification factor (Pearson *r* = 0.7, *P* = 0.056). Dashed line represents the 95 % confidence interval of the control data. e. Maximum decoder accuracy per animal, as a function of DA reward amplification factor. The decoder was trained using 50 neurons to discriminate R from R+L trials. (Pearson *r* = 0.6, *P* = 0.08). Dashed line represents the 95 % confidence interval of the control data. f. Mean firing rate of two MSNs showing an excitatory (top) and inhibitory (bottom) response to DA neuron activation. Data are from the supra-reward condition (animals exhibiting over 80 % decoding accuracy in panel e). Data represent mean ± SEM. g. Top: Selectivity index of 423 neurons pooled across 4 Chrimson-expressing mice. Bottom: Percentage of neurons that were significantly selective for R versus R+L trials, as a function of time. Data are from the supra-reward condition (animals exhibiting over 80 % decoding accuracy in panel e).

In order to examine DA’s modulatory effects during non-rewarding stimuli, in a subset of animals we included a block of trials consisting of randomly interleaved neutral auditory tone and laser-paired tone stimuli. As before, neural activity could only reliably distinguish these conditions when DA signals were amplified above tone-matched levels (**Supplemental Fig. 6**). Together, the DA activation experiments suggest that only supra-reward DA signals are capable of producing consistent changes in either spontaneous, reward-evoked, or auditory stimulus-evoked spiking in the ventral striatum.

## Discussion

This study sought to test and clarify the longstanding prediction that striatal spiking activity is strongly and rapidly regulated by DA neuron signals. This phenomenon is a linchpin in the popular model that DA plays a major role not only in shaping future behavior (i.e., via learning-dependent plasticity mechanisms), but in imminent or ongoing actions ^4^. While the impact of DA and DA neurons on striatal excitability and spiking has been demonstrated numerous times, previous approaches involved DA manipulations that lacked either subsecond temporal precision or calibration to a physiologically relevant stimulus such as reward. This study addressed prior limitations by taking advantage of optogenetics to transiently perturb DA neurons, together with a dual electrophysiological and DA recording approach, which enabled a systematic analysis of striatal dynamics under various calibrated levels of DA. The study reveals two main findings that challenge the above prediction. First, that DA signals at typical levels encountered in research on natural rewards are neither necessary nor sufficient to cause consistent, robust changes in spontaneous or reward-evoked ventral striatal neuronal firing. This indicates that reward-related striatal spiking activity is largely uncoupled from rapid increases in DA signaling that accompany unexpected reward events. These results suggest that on short timescales striatal reward activity is primarily driven by non-DA inputs, including cortical and thalamic glutamatergic projections ^37^. The second major finding is that DA neurons are capable of producing strong and rapid changes in striatal spiking activity under artificial, supra-reward DA signaling conditions. The transition from small to large electrophysiological effects became gradually more apparent with increasing DA levels. To estimate this transition point, we can refer to the population decoding analysis, which provided a sensitive method for detecting changes in neural activity. Interpolating these results, the transition to the high decoding accuracy (80 %) regime occurred when DA signals were amplified around three to four times above reward-matched levels (**Fig. 5g, 6e**).

A major question is whether this work can be reconciled with a sizable body of evidence that appears to contradict our findings either directly via electrophysiological measurements ^7, 8^, or indirectly via behavioral measurements ^9–11^. On one hand, a subset of those experiments may have inadvertently over-stimulated DA neuron cell bodies or terminals, causing supra-physiological neurotransmitter release in the striatum. There is further support for this possibility from a study showing that animal motion effects are only evident when substantia nigra (SNc) DA levels are raised five times above reward-matched levels ^19^. This five-fold behavioral threshold is remarkably similar to our electrophysiologically determined threshold estimate, suggesting the same underlying modulatory mechanisms. While it is tempting to speculate on the prevalence of engaging supra-physiological mechanisms within the literature, it is noted that since optogenetically evoked DA signals are often not measured let alone calibrated, this possibility can neither be firmly proved nor disproved. On the other hand, a number of studies used either DA neuron inhibition or a pre-calibrated level of activation, thereby mitigating the risk to engage supra-physiological DA release mechanisms. In one such experiment, optogenetically manipulating VTA DA neurons bidirectionally altered the probability of initiating reward-motivated approach behavior ^10^. Two separate studies showed that SNc DA manipulations biased the probability of selecting a particular action over another ^13, 14^. Elsewhere, optogenetic SNc DA manipulations modulated the probability of initiating an action at a given moment in time ^12^. Moreover, an earlier study from our group showed that inhibiting SNc DA neurons reduced the probability of initiating reward-anticipatory movements ^20^. These statistically significant behavioral changes may appear to contradict the findings in this work, but a crucial observation is that the average change in movement or action probability seen in those studies is generally small, particularly in the case of DA inhibition. By comparison, our neural decoding analysis suggests a downstream area can theoretically distinguish reward-evoked striatal population dynamics with or without DA signals in only 8 % more trials relative to controls (**Fig. 3b**). Thus, when considering inhibitory manipulations, the size of previously reported behavioral effects, and the upper bound on the size of electrophysiological effects shown here, are comparable. The most parsimonious, unified interpretation of these observations is that DA neurons modulate striatal activity, and consequently behavior, in a weak probabilistic manner, i.e., on a small minority of trials. Our results place important physiological constraints on the contribution of DA neurons to striatal dynamics. The probabilistic effect on striatal dynamics is normally small in size (with spiking changes appearing on the order of 10 % of trials or less), but is significantly enhanced by non-physiological DA signals.

Multiple synaptic and microcircuit-level mechanisms could potentially explain how DA neurons modulate striatal spiking activity on subsecond timescales ^1^. Disentangling these contributions is the subject of other work and was not the purpose of this study, but we can speculate on potential mechanisms. Previous work has shown that stimulating DA terminals can rapidly enhance the firing of D1 receptor (D1R) expressing MSNs via a D1R-dependent mechanism ^7^. DA also modulates lateral inhibition between MSNs, which may alter spiking activity ^38^. In addition to DA itself, there may be a role for DA neuron co-release of glutamate and GABA ^39, 40^, as both of these neurotransmitters are well suited to drive rapid changes in striatal firing. There is evidence that blocking glutamate co-release attenuates DA stimulation-evoked spiking in the ventral striatum ^8^. However, since these mechanistic studies did not appear to discriminate between reward-matched and supra-reward (or physiological and supra-physiological) DA signals, it remains unclear to what extent the same mechanisms were involved in our experiments.

A further open question is whether the electrophysiological findings generalize across different behaviors and striatal subregions. First, we cannot directly rule out the presence of stronger modulatory effects during other types of tasks. Nevertheless, we examined rapid DA modulation of two highly distinct modes of striatal activity – spontaneous and reward-evoked firing. Since similar trends were observed under these conditions as well as during neutral auditory stimuli, it is unclear what mechanisms could cause a dramatically higher effect size under different tasks. However, this assumption may not hold for disorders involving altered DA signaling such as addiction. In any event, the recording methods presented here provide a path to resolving the generality question directly. Second, our experiments were restricted to measuring ventral striatal dynamics during somatic VTA DA manipulation. One study suggested that DA has a stronger effect on synaptic transmission in the ventral compared to the dorsal striatum ^41^. There is evidence that DA glutamate co-release mechanisms may be biased toward the nucleus accumbens ^39^. Additionally, DA drives heterogeneous synaptic responses on cholinergic interneurons in different striatal subregions ^36^. It is therefore conceivable that DA neurons may differentially modulate dorsal and ventral striatal dynamics on short timescales. But there is no known mechanism that would suggest these effects are enhanced in the dorsal striatum. A related issue is the specificity of our DA neuron manipulations to the striatum. In order to maximize the likelihood of altering neural dynamics, we intentionally targeted all types of VTA DA neurons instead of only the population projecting to the ventral striatum. Many of the aforementioned behavioral studies also did not exclusively target striatal projecting DA neurons. This raises the potential pitfall of off-target network effects of opto-stimulation ^42^. For example, under certain stimulation conditions VTA DA neurons can rapidly alter the dynamics of prefrontal cortical neurons ^43, 44^, which in turn project to the striatum. However, even if such indirect effects are present, then the direct contribution of DA neurons to striatal dynamics would likely be smaller than our current approach suggests, thus further strengthening the main conclusions of this work.

Taken together, our results emphasize the need to carefully distinguish between the modulatory function that DA neurons normally have in shaping ventral striatal dynamics, and the role they are capable of serving under potentially non-physiological conditions. One possible approach to circumvent this issue is to measure and calibrate DA stimulation levels with respect to a known reference signal on a per animal basis ^10, 12, 19^. However, we suggest that in order to avoid potential calibration errors, it is essential to include experiments that assess in vivo DA function through inhibitory manipulations. Here, transiently inhibiting DA neurons produced only small changes in spontaneous and reward-evoked striatal spiking. Thus, the view that DA plays a major role in rapidly influencing behavior by modulating striatal activity on subsecond timescales, may need to be reevaluated. This would have profound implications for understanding the apparent duality of DA neuron function in learning and online motor control.

## Methods

### Animals

DAT-Cre mice (Strain # 006660, Jackson Laboratories) and VGAT-Cre mice (Strain # 028862, Jackson Laboratories), 8-18 weeks old at the time of the first surgery, were used for experiments involving optogenetic manipulation of dopaminergic neurons or GABAergic neurons, respectively. Both male and female mice were used, and the mice were maintained as heterozygotes on a C57BL/6J background. Animals were kept on a 12 hour light/dark cycle, and were group housed until the first surgery. All procedures were approved by the University of California, Los Angeles Chancellor’s Animal Research Committee.

### Surgical procedures

Surgery was performed under isoflurane anesthesia and aseptic conditions using a stereotaxic apparatus (Model 1900, Kopf Instruments). The procedure consisted of three principal components: injection of adeno-associated virus (AAV) into the ventral tegmental area (VTA) and ventral striatum, placement of a fiber-optic implant in the VTA, and attachment of stainless-steel head fixation bars to the skull. AAV for optogenetics (UNC Vector Core) was unilaterally injected into the VTA, consisting of 500 nl of Cre-dependent AAV expressing either AAV5/EF1a-DIO-eNpHR3.0-mCherry, AAV5/Syn-Flex-ChrimsonR-tdTomato, or AAV5/EF1a-DIO-mCherry (coordinates relative to bregma: 3.30 mm posterior, 0.50 mm lateral, 4.30 mm ventral). AAV expressing the fluorescent DA sensor dLight1.2 under the synapsin promoter (pAAV-hSyn-dLight1.2) was also unilaterally injected into the striatum (coordinates relative to bregma: 1.30 mm anterior, 1.25 mm lateral, 4.25 mm ventral). pAAV-hSyn-dLight1.2 was a gift from Lin Tian (Addgene plasmid # 111068) ^34^. Following viral injection, a ferrule-coupled optical fiber implant (0.2 mm diameter, 0.22 NA, Thor Labs) was placed with the tip approximately 0.2 mm above the viral injection site. After surgery, all animals were individually housed and were given daily carprofen injections (5mg/kg, subcutaneous) for the first three post-operative days. Over the first post-operative week, animals were administered ibuprofen and amoxicillin dissolved in their drinking water. The mice were given a recovery period of at least two weeks prior to behavioral training.

### Behavioral task

Mice were food restricted but given ad libitum access to water to maintain around 90 % of their baseline weight. Animals were habituated to head fixation and trained to reliably consume uncued rewards (6 μl, 10 % sweetened condensed milk) delivered via an audible solenoid valve. Rewards were delivered via a tube within an infrared lick detection port located approximated 5 mm from the mouth. Mice had to extend their tongue out of the mouth multiple times to consume the reward and these tongue extensions were detected as licks. Mean lick rates were calculated based on the average number of consummatory licks from 0-1 s following reward delivery. During behavioral training, mice were given rewards (20-30 s inter-trial interval, ITI, 100 trials per day). Habituation to head fixation and lick training took place over the course of 7 total days. A subset of animals was also habituated to an audible tone on the last two days of training (12 kHz tone, 0.5 s duration, 20-30 s ITI, 100 trials per day). After the training period, mice then proceeded to the experiment involving electrophysiology, photometry, and optogenetics.

### Optogenetics

Optogenetic manipulations were organized into two principal sections, with the first block consisting of reward trials and laser-paired reward trials (R and R+L), and the second block consisting of unpaired laser stimulation (589 nm, 5 mW power, MGL-F-589-100 mW, CNI Laser). For eNpHR3-mediated opto-inhibition, laser stimulation was applied continuously for 0.5 s. The first block consisted of 40 R and 40 R+L trials randomly interleaved (20-30 s ITI). The second block consisted of 80-100 trials where laser was delivered alone (20-30 s ITI); in order to maintain a consistent number of trials across different analyses, only the last 40 trials were analyzed. For experiments involving Chrimson-mediated opto-activation, laser stimulation was pulsed at either 4, 20, or 40 Hz over a 0.5 s period (10 ms pulse width). The first block was similar to the inhibition experiments consisting of 38-40 each of R and R+L trials (laser was pulsed at 40 Hz). The second block consisted of 40 trials each of 4, 20, and 40 Hz unpaired laser stimulation presented in random order. One animal did not undergo 4 Hz laser stimulation. A subset of the animals used for the activation experiments also completed an additional block where an auditory tone was presented with half the trials randomly paired with optogenetic stimulation (40 T and 40 T+L trials).

### Electrophysiology

To measure reward- and optogenetically evoked rapid changes in DA signaling alongside neural spiking responses in the ventral striatum, an opto-probe containing a 256 electrode silicon microprobe (type 256AS, developed by our lab ^45^) attached to a low autofluorescence photometry fiber positioned about 0.15 mm above the nearest electrode recording sites (0.43 mm diameter, 0.48 NA, Doric Lenses). The silicon microprobe contained four shanks separated by 0.4 mm. Each shank contained 64 electrodes spanning a total length of 1.05 mm. This device allowed recordings from a wide area of the nucleus accumbens in the ventral striatum. Prior to electrophysiological recordings, a second surgery was performed under isoflurane anesthesia in which a rectangular craniotomy over the striatum was opened and then covered with a silicone sealant (Kwik-Cast, World Precision Instruments). The animal was allowed to recover for at least 3 hours before inserting the opto-probe into the nucleus accumbens region of the ventral striatum (ventral tip of probe shanks: 1.30 mm anterior, 0.65-1.85 mm lateral, 5.1 mm ventral). The shanks of the opto-probe were coated with a fluorescent dye prior to insertion (DiI or DiD, Thermo Fisher Scientific). Electrophysiological recordings were conducted at a sampling rate of 25 kHz per electrode using a commercial multichannel data acquisition (DAQ) system (C3316 and C3004, Intan Technologies). Spike sorting was carried out with Kilosort ^46^, using bandpass filtered data (3 pole Butterworth filter with a passband of 0.6-7 kHz). After identifying single-units, raw data were refiltered with a wider passband (0.3-7 kHz) and then classified as putative MSNs, FSIs, and TANs. The classification procedure was based on spike trough-to-peak duration and baseline firing rate ^47^. Putative FSIs were defined by a narrow spike waveform (0.45 ms maximum duration). MSNs and TANs were both defined by wider waveforms (0.475-2 ms duration). TANs were separated from MSNs by the regularity of their baseline firing (maximum TAN coefficient of variation: 1). The minimum baseline firing was defined as 0.01 Hz for MSNs and FSIs and 2.5 Hz for TANs, and the maximum was defined as 20 Hz for MSNs and TANs. Unless specified, all analysis of neural activity used all cell types including unclassified units, in order to avoid possible bias arising from cell-type specific modulatory effects as well as classification errors. For assessing changes in neural activity (see the Selectivity Index and Population decoding sections below) firing rates were calculated by binning spikes in 50 ms increments using a three bin sliding window average.

### Fiber photometry

Photometry of dLight fluorescence was conducted synchronously with electrophysiological recordings using the opto-probe described above. The photometry fiber was coupled via a fiber patch cord to a four-port connectorized fluorescence mini cube (FMC4_AE(405)_E(460-490)_F(500-550)_S, Doric Lenses), with one excitation port (460-490 nm) used for dLight1.2 fluorescence, and a 500-550 nm detection port. Fluorescence excitation light was delivered via a 465 nm light emitting diode oscillated at 211 Hz. The emitted signal was then sent to a low noise femtowatt photoreceiver (Model 2151, Newport) connected to a lock-in amplifier (SR810, Stanford Research Systems). The demodulated signal was sampled at 25 kHz by a DAQ (Intan Technologies), and then downsampled to 1 kHz for storage. Offline analysis involved downsampling the signal again to 20 Hz. The fractional change in fluorescence (ΔF/F) was calculated with respect to the average baseline signal 1 s prior to the onset of a trial. The average fractional change in fluorescence was determined by calculating the average value of ΔF/F for a given trial condition from 0-1 s post-stimulus onset. The factor increase in DA was calculated as the ratio of the maximum laser-evoked ΔF/F to the maximum reward-evoked ΔF/F signal. The DA reward amplification factor was calculated as the ratio of the maximum ΔF/F on R+L trials to the maximum ΔF/F on R trials.

### Photometry depth test

To determine whether the estimated factor increase in DA was consistent across different depths spanned by the electrode array, a group of DAT-Cre mice were injected with AAV5/Syn-Flex-ChrimsonR-tdTomato in the VTA and pAAV-hSyn-dLight1.2 in the ventral striatum in a procedure identical to that described in the Surgical procedures section. The mouse also underwent identical behavioral training and craniotomy procedures. However, for recording, instead of using an opto-probe, a photometry fiber was lowered to a site corresponding to the top of the area targeted by the electrodes (1.30 mm anterior, 1.25 mm lateral, 4.1 mm ventral). It was then lowered in 0.2 mm increments to target five additional depths (maximum depth: 5.1 mm). At each depth, the mouse was given a randomly interleaved combination of 10 reward trials and 10 trials of 40 Hz pulsed optogenetic stimulation. Between each depth measurement, a 10 minute waiting period ensued for tissue surrounding the optical fiber to stabilize. Before each recording commenced, mice were given 2-3 rewards to ensure that licking behavior was maintained.

### Immunohistochemistry

Mice were anesthetized with isoflurane and transcardially perfused with phosbate buffered saline followed by 10 % neutral buffered formalin. Brains were stored in the neutral buffered formalin solution for at least 1 day at 4 °C before being sectioned at a thickness of 100 μm on a vibratome. Sections were blocked using donkey serum before antibody incubations. VTA slices were incubated with rabbit anti-dsRed polyclonal antibody (632496, Takara) as the primary antibody (1:1000 dilution) overnight at 4 °C. Striatal slices were incubated with chicken anti-GFP (ab13970, Abcam) as the primary antibody (1:1000 dilution) overnight at 4°C. VTA slices were then incubated with Alexa Fluor 647-conjugated donkey antibody to rabbit IgG (711-605-152, Jackson ImmunoResearch) as the secondary antibody (1:200 dilution) for 4 hours. Striatal slices were incubated with Alexa Fluor 488-conjugated donkey antibody to chicken IgG (703-545-155, Jackson ImmunoResearch) as the secondary antibody (1:200 dilution) for 4 hours. Sections were then mounted using tissue-mounting medium and imaged under a confocal or epifluorescence microscope.

### Selectivity index

Single-neuron selectivity indices were used in two different ways. The first selectivity index was used to determine a neuron’s discrimination of R+L versus R trials, or T+L versus T trials. The second selectivity index was used to determine a neuron’s discrimination of unpaired reward or laser trials versus a preceding baseline period. In both cases the selectivity index was calculated by subtracting 0.5 from the area under the ROC curve and then multiplying the result by 2. The selectivity index was calculated separately for each time bin from −1 to 3 s post-stimulus. The significance level of neural responses was assessed by comparing the observed difference in mean to the resampled difference in mean using randomly shuffled trial assignments (400 permutations). A neuron was defined as being significantly modulated by a stimulus, or selective between two stimulus conditions, if three or more consecutive time bins from 0-1 s showed a statistically significant change in firing (observed difference in mean was greater than 95 % of resampled differences in mean).

### Population decoding

As with the selectivity index described above, population decoding was used in two different ways. The first type of decoder was trained to distinguish population dynamics on R+L versus R trials, or T+L versus T trials. The second type of decoder was trained to distinguish population dynamics on unpaired laser trials versus a preceding baseline period. For analysis of DA neuron opto-inhibition and GABA neuron opto-activation effects, the decoder was trained on data pooled across all animals, with the number of neurons varied from 5 to 300. For analysis of DA neuron opto-activation effects, the decoder was trained on 50 simultaneously recorded neurons. The decoder was based on a linear support vector machine (SVM) learning algorithm ^48^, and utilized 80 % of trials for training and the remaining 20 % for testing the accuracy. The mean decoder accuracy was calculated from the average performance of 50 random neuron selections, and 100 random train/test trial selections. The decoder was separately trained using randomly shuffled trial assignments. The 95 % confidence interval was calculated from testing decoder performance on these shuffled datasets ^49^.

### Statistics

Statistical analysis utilized standard functions in Matlab (MathWorks) and Prism (GraphPad Software). Data collection and analysis were not performed with blinding to the experiments. No statistical methods were used to predetermine sample sizes, but our sample sizes are similar to those used in previous publications. The sample size, type of statistical test, and P values are indicated in figure legends. Data distribution was assumed to be normal, but not specifically tested. T-tests were two sided. One-way ordinary or RM ANOVA was followed by Tukey’s post hoc test for multiple comparisons. Two-way RM ANOVA was followed by Sidak’s post-hoc test for multiple comparisons.

## Data and Code availability

The data and Matlab code that support the findings of this study are available from the corresponding author upon request.

## Acknowledgments

SCM was supported by NIH grants NS100050, NS125877, DA042739, DA005010, and NSF NeuroNex Award 1707408.

## Author contributions

Study design: CL, KL, SCM; data collection: CL, KL, AKW, TD; analysis and interpretation of results: CL, KL, LY, SCM; manuscript preparation: CL, KL, SCM.

## Supplemental Figures

**Supplemental Fig. 1:**
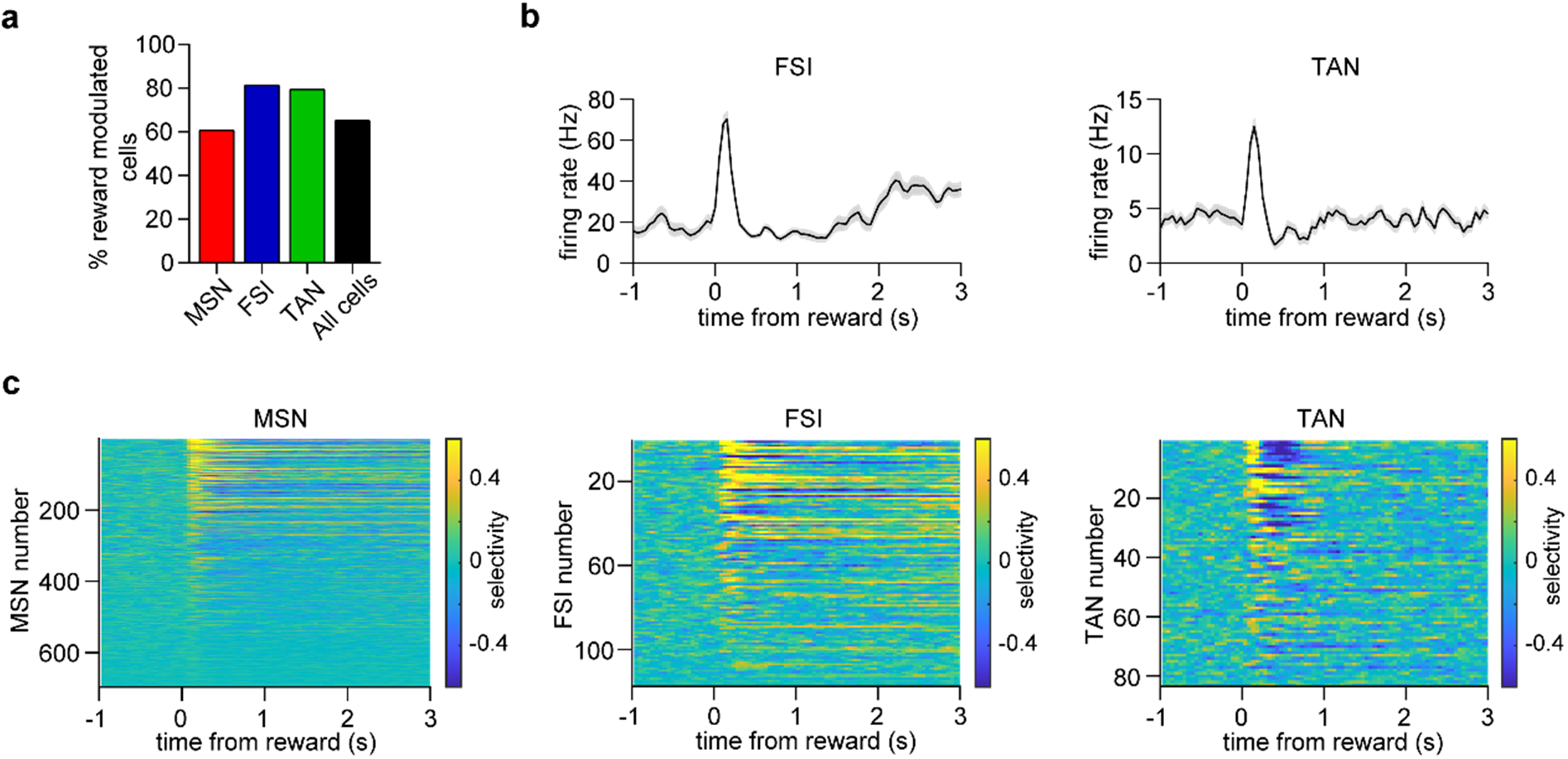
Reward response of electrophysiologically classified cell types in the ventral striatum. a. Percentage of each cell type that showed a significant reward response. b. Mean firing rate of one putative FSI (left) and TAN (right) unit during reward delivery. c. Selectivity index of a population of 697 MSNs, 118 FSIs, and 83 TANs during reward delivery. Selectivity of neural activity is calculated relative to baseline. All data in this figure represent mean ± SEM.

**Supplemental Fig. 2:**
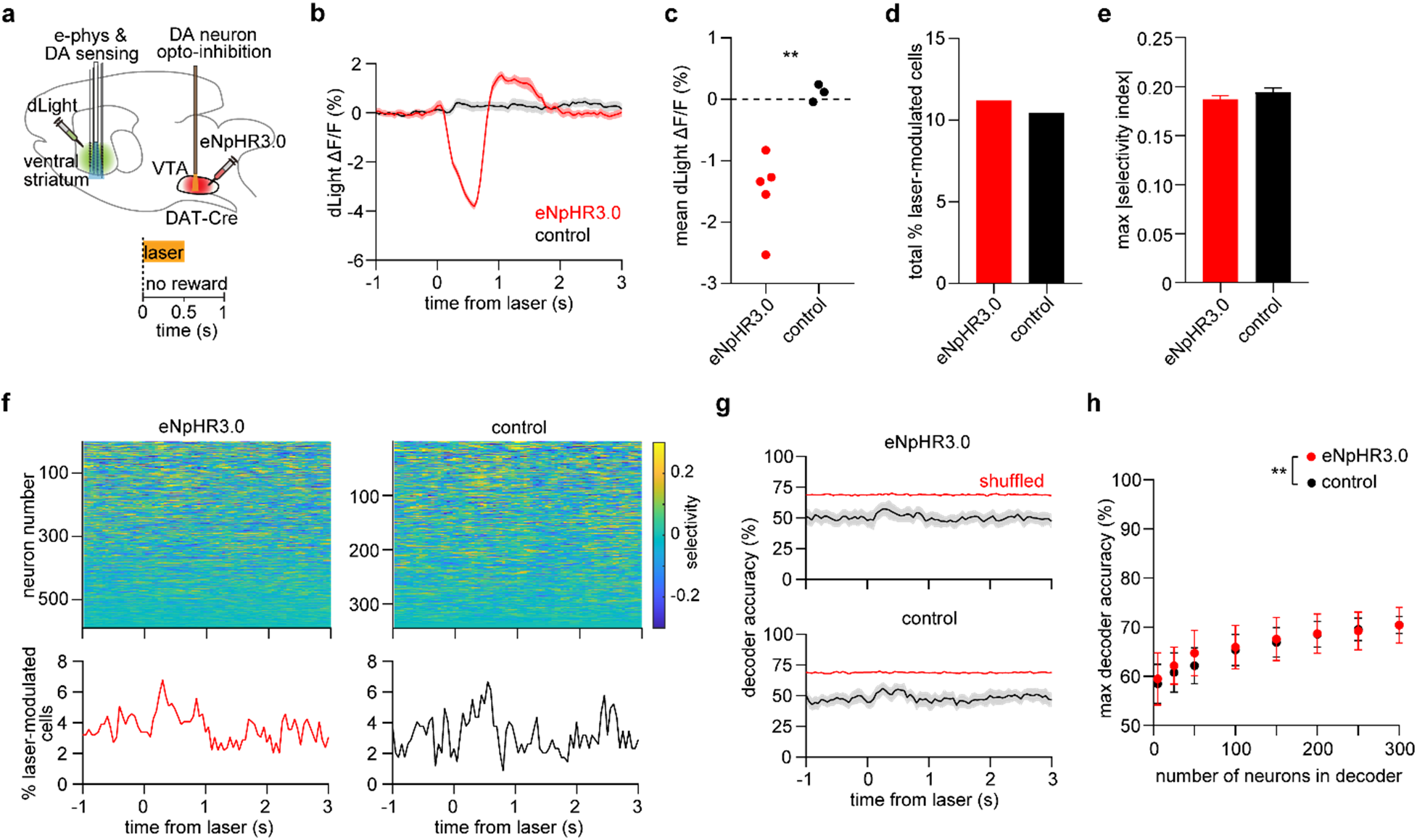
Small effect of inhibiting VTA DA neurons on spontaneous striatal spiking activity. a. Top: Experimental approach. Bottom: task schematic in which 0.5 s laser stimulation is delivered in the absence of reward or other stimuli. b. dLight fractional fluorescence change signal from one eNpHR3.0-expressing mouse (red) and one opsin-free control mouse (black). c. Mean laser-evoked dLight fluorescence (n = 5 eNpHR3.0 and 3 control mice, unpaired t-test, *t* = 4.2, \*\**P* = 0.005). d. Total percentage of neurons that were significantly modulated by the laser relative to baseline (n = 66 out of 589 cells in the eNpHR3.0 group, n = 36 out of 344 cells in the control group, chi square test for proportions, *χ^2^* = 0.2, *P* = 0.8). e. Maximum selectivity index per neuron (n = 589 cells in the eNpHR3.0 group, n = 344 cells in the control group, unpaired t-test, *t* = 1.1, *P* = 0.3). Selectivity of neural activity is calculated relative to baseline. Data represent mean ± SEM. f. Top: Selectivity index of 589 neurons pooled across 5 eNpHR3.0-expressing mice, and 344 neurons pooled across 3 control mice. Bottom: Percentage of neurons that were significantly modulated by the laser relative to baseline, as a function of time. g. Mean accuracy of an SVM decoder trained using 50 neurons to discriminate laser-evoked from baseline activity. Red line indicates the 95 % confidence interval of decoder performance trained on trial-shuffled data. Top: neurons selected from the eNpHR3.0 group. Bottom: neurons selected from the control group. Shaded area represents the SD across 50 random drawings of neurons. h. Maximum decoder accuracy as a function of neuron number (two-way ANOVA, group effect: *F_1,784_* = 8, *P* = 0.004, neuron number effect: *F_7,784_* = 116, *P* < 0.0001). Post-hoc Sidak’s test: \*\**P* = 0.005 for n = 50 neurons; all other comparisons between the eNpHR3.0 and control group were not significant. Data represent the mean and SD across 50 random drawings of neurons.

**Supplemental Fig. 3:**
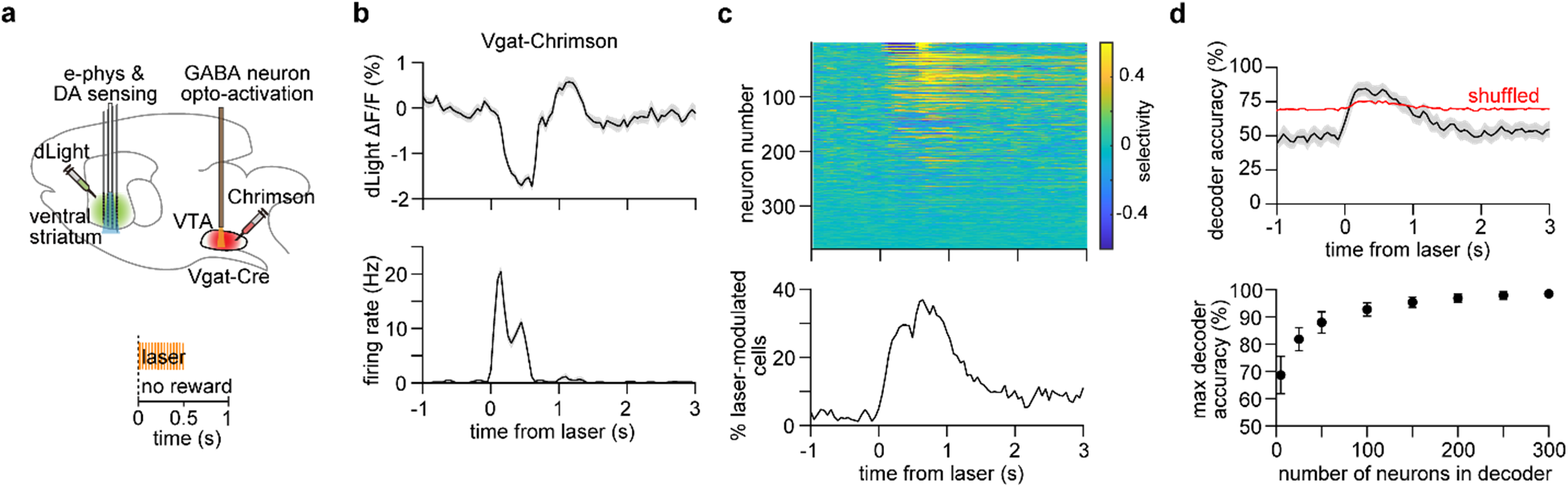
Strong effect of activating VTA GABAergic neurons on spontaneous striatal spiking activity. a. Top: Experimental approach. Bottom: task schematic in which 0.5 s pulsed laser stimulation is delivered in the absence of reward or other stimuli. b. Top: dLight fractional fluorescence change signal during laser stimulation from one Vgat-Chrimson mouse. Bottom: Mean firing rate of one neuron. Data represent mean ± SEM. c. Top: Selectivity index of 379 neurons pooled across 3 Vgat-Chrimson mice. Bottom: Percentage of neurons that were significantly modulated by the laser relative to baseline, as a function of time. d. Top: Mean accuracy of an SVM decoder trained using 50 neurons to discriminate laser-evoked from baseline activity. Shaded area represents the SD across 50 random drawings of neurons. Red line indicates the 95 % confidence interval of decoder performance trained on trial-shuffled data. Bottom: Maximum decoder accuracy as a function of neuron number (one-way ANOVA, neuron number effect: *F_7,392_* = 441, *P* < 0.0001). Post-hoc Sidak’s test: *P* < 0.0001. Data represent the mean and SD across 50 random drawings of neurons.

**Supplemental Fig. 4:**
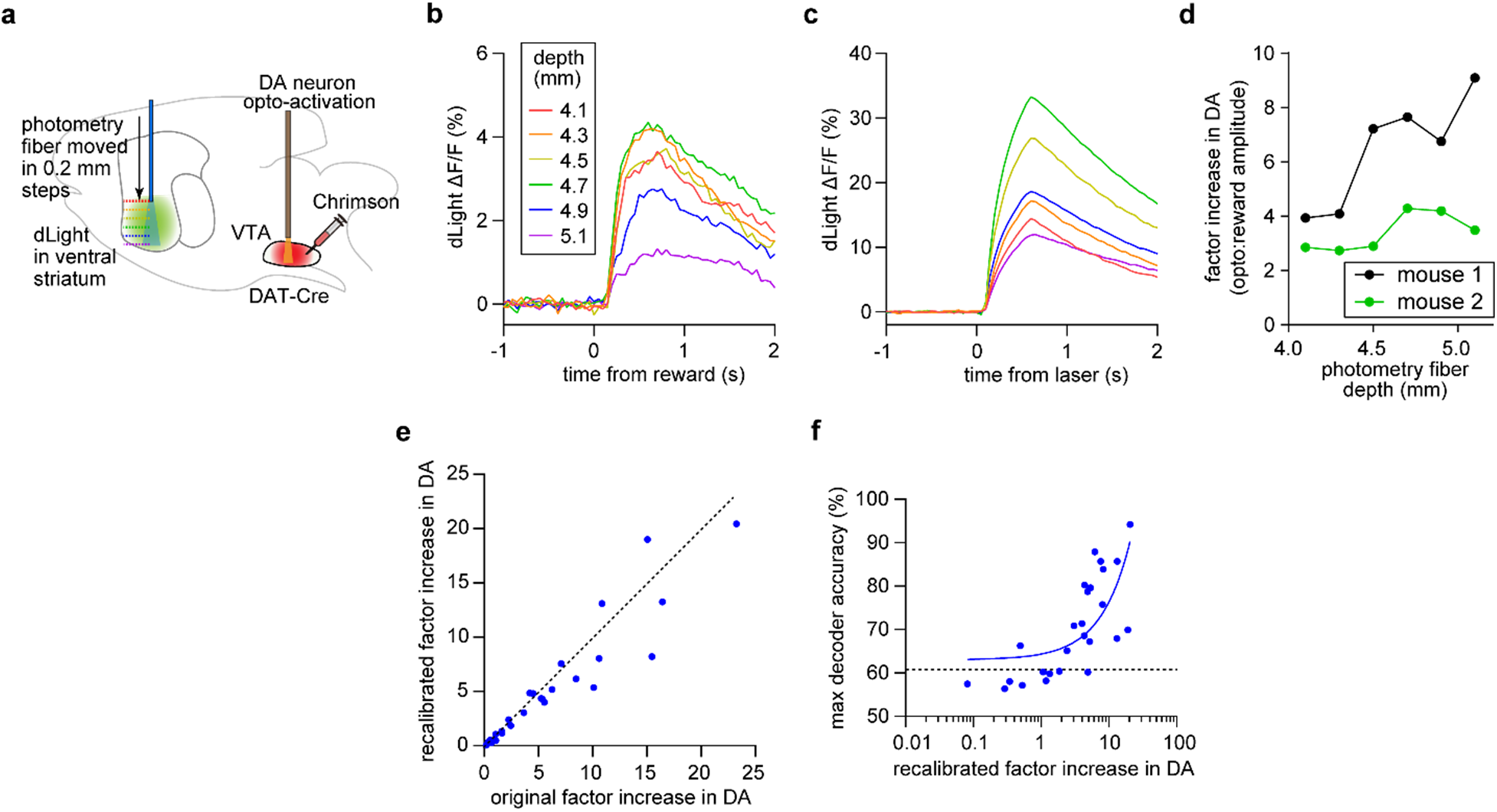
Assessing sources of variability in estimating the factor increase in DA. a. Experimental approach of the dLight photometry depth test. b. Reward-evoked dLight fractional fluorescence change signal from one Chrimson-expressing mouse. The plots are color-coded by photometry fiber depth from bregma. c. Laser-evoked dLight fractional fluorescence change signal from one Chrimson-expressing mouse. The plots are color-coded by photometry fiber depth from bregma. d. Factor increase in DA (the ration of laser-evoked to reward-evoked dLight signal amplitudes) calculated from data at different photometry fiber depths. Data are from n = 2 mice. e. Recalibrated versus original factor increase in DA. The recalibrated factor used only the first 10 % of reward trials (n = 26 sessions from 9 Chrimson-expressing mice, paired t-test, *t* = 2.1, *P* = 0.04). f. Maximum decoder accuracy per recording session, as a function of recalibrated factor increase in DA. The decoder was trained using 50 neurons to discriminate laser-evoked from baseline activity. Chrimson group data are positively correlated with the factor increase in DA (Pearson *r* = 0.9, *P* < 0.0001). Dashed line represents the 95 % confidence interval of the control data. Blue line represents the linear fit to the Chrimson data.

**Supplemental Fig. 5:**
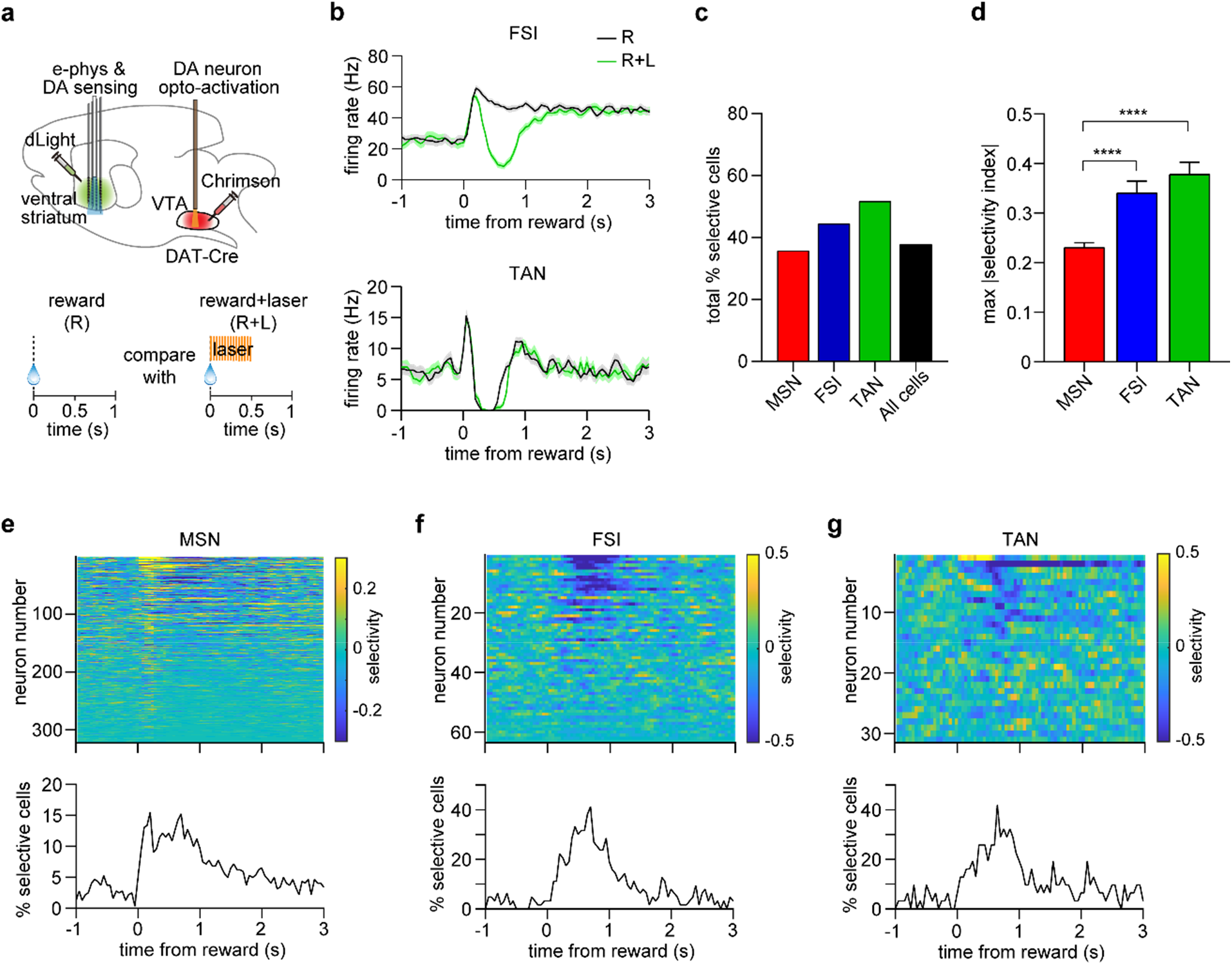
Effect of supra-reward DA signals on reward response of electrophysiologically classified cell types in the ventral striatum. a. Top: Experimental approach. Bottom: task schematic in which reward trials (R) are compared to laser-paired reward trials (R+L). All data in this figure are from the supra-reward condition (4 mice exhibiting over 80 % decoding accuracy in Fig. 6e). b. Mean firing rate of a putative FSI (top) and TAN (bottom) MSNs on R and R+L trials. Note the broadening of the TAN post-reward pause response. Data represent mean ± SEM. c. Total percentage of each cell type that was selective for R versus R+L trials. d. Mean of the maximum absolute value of the selectivity index per neuron, grouped by cell type (n = 322 MSN, 63 FSI, 31 TAN, one-way ANOVA, cell type effect: *F_2,413_* = 18, *P* < 0.0001). Post-hoc Sidak’s test: \*\*\*\**P* < 0.0001. Data represent mean ± SEM. e. Top: Selectivity index of 322 MSNs pooled across 4 Chrimson-expressing mice. Bottom: Percentage of neurons that were significantly selective for R versus R+L trials, as a function of time. f. Top: Selectivity index of 63 FSIs pooled across 4 Chrimson-expressing mice. Bottom: Percentage of neurons that were significantly selective for R versus R+L trials, as a function of time. g. Top: Selectivity index of 31 TANs pooled across 4 Chrimson-expressing mice. Bottom: Percentage of neurons that were significantly selective for R versus R+L trials, as a function of time.

**Supplemental Fig. 6:**
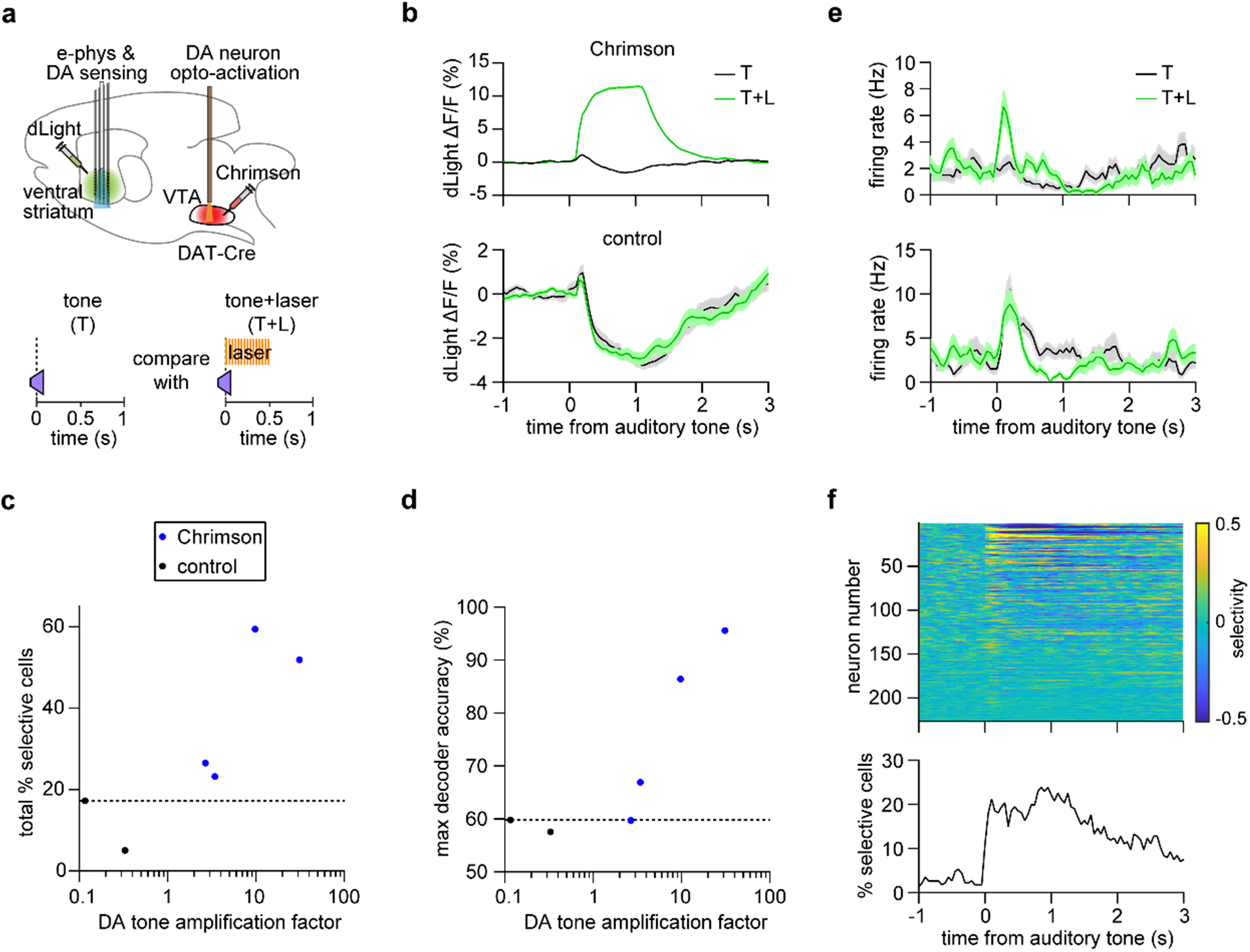
Effect of activating DA neurons during auditory stimulus processing in the ventral striatum. a. Top: Experimental approach. Bottom: task schematic in which neutral auditory tone trials (T) are compared to laser-paired tone trials (T+L). b. dLight fractional fluorescence change signal from one Chrimson-expressing mouse (top) and one opsin-free control mouse (bottom) on tone (T) and laser-paired tone (T+L) trials. Data represent mean ± SEM. c. Percentage of neurons per animal that were selective for T versus T+L trials, as a function of DA tone amplification factor (an amplification factor of one corresponds to tone-matched DA levels). n = 4 Chrimson-expressing and 2 control mice (Pearson *r* = 0.6, *P* = 0.37). Dashed line represents the 95 % confidence interval of the control data. d. Maximum decoder accuracy per animal, as a function of DA tone amplification factor. The decoder was trained using 50 neurons to discriminate R from R+L trials. n = 4 Chrimson-expressing and 2 control mice (Pearson *r* = 0.9, *P* = 0.12). Dashed line represents the 95 % confidence interval of the control data. e. Mean firing rate of two putative MSNs on T and T+L trials. Data are from the supra-reward condition (mice exhibiting over 80 % decoding accuracy in panel d). Data represent mean ± SEM. f. Top: Selectivity index of 226 neurons pooled across 2 Chrimson-expressing mice. Bottom: Percentage of neurons that were significantly selective for T versus T+L trials, as a function of time. Data are from the supra-reward condition (mice exhibiting over 80 % decoding accuracy in panel d).

